# Progression of regional grey matter atrophy in multiple sclerosis

**DOI:** 10.1101/190116

**Authors:** Arman Eshaghi, Razvan V. Marinescu, Alexandra L. Young, Nicholas C. Firth, Ferran Prados, M Jorge Cardoso, Carmen Tur, Floriana De Angelis, Niamh Cawley, Wallace Brownlee, Nicola De Stefano, M. Laura Stromillo, Marco Battaglini, Serena Ruggieri, Claudio Gasperini, Massimo Filippi, Maria A. Rocca, Alex Rovira, Jaume Sastre-Garriga, Jeroen Geurts, Hugo Vrenken, Viktor Wottschel, Cyra E Leurs, Bernard Uitdehaag, Lukas Pirpamer, Christian Enzinger, Sebastien Ourselin, Claudia A.G. Wheeler-Kingshott, Declan Chard, Alan J. Thompson, Frederik Barkhof, Daniel C. Alexander, Olga Ciccarelli, on behalf of the MAGNIMS study group

## Abstract

Grey matter atrophy is present from the earliest clinical stages of multiple sclerosis (MS), but the temporal ordering is poorly understood. We aimed to determine the sequence in which grey matter regions become atrophic in MS, and its association with disability accumulation.

In this longitudinal study, we included 1,417 subjects: 253 with clinically-isolated syndrome (CIS), 708 relapsing-remitting MS (RRMS), 128 secondary-progressive MS (SPMS), 125 primary-progressive MS (PPMS), and 203 healthy controls from 7 European centres. Subjects underwent repeated MRI scanning (total number of scans 3,604); the mean follow-up for patients was 2.41yrs (SD±1.97). Disability was scored using the Expanded Disability Status Scale (EDSS). We calculated the volume of brain grey matter regions and brainstem using an unbiased within-subject template. We used an established data-driven event-based model (EBM) to determine the sequence of occurrence of atrophy and its uncertainty. We assigned each subject to a specific EBM stage, based on the number of their atrophic regions. We used nested linear mixed-effects regression models to explore the associations between the rate of increase in the EBM stages over time, disease duration and annual rate of EDSS gain.

The first regions to become atrophic in CIS and relapse-onset MS patients (RRMS and SPMS) were the posterior cingulate cortex and precuneus, followed by the middle cingulate cortex, brainstem and thalamus. The sequence of atrophy in PPMS showed a similar involvement of the thalamus, cuneus, precuneus, and pallidum, followed by the brainstem and posterior cingulate cortex. The cerebellum, caudate and putamen showed early atrophy in relapse-onset MS and late atrophy in PPMS. Patients with SPMS showed the highest EBM stages (highest number of atrophic regions, all *p*<0.001) at study entry. Rates of increase in EBM stages were significantly different from healthy controls in all MS phenotypes, except for CIS. The increase in the number of atrophic regions (EBM stage) was associated with disease duration in all patients. EBM stage was associated with disability accumulation in RRMS independent of disease duration (*p*<0.0001).

This data-driven staging of atrophy progression in a large MS sample demonstrates that grey matter atrophy spreads to involve more regions over time. The sequence in which regions become atrophic is reasonably consistent across MS phenotypes. The spread of atrophy was associated with disease duration, and disability accumulation in RRMS.

**Abbreviations:**

MS: multiple sclerosis
GM: grey matter
FLAIR: Fluid Attenuated Inversion Recovery
PPMS: primary progressive multiple sclerosis; primary-progressive MS

## Introduction

Multiple sclerosis (MS) is an inflammatory demyelinating disease of the central nervous system with a prominent neurodegenerative component. Brain atrophy, as assessed by MRI, develops at a faster rate in people with MS than healthy controls. Whole brain atrophy is the result of grey matter and to a lesser extent of white matter atrophy (Fisher *et al*., 2008), which is related to long term disability in MS (Fisniku *et al*., 2008; Filippi *et al*., 2013). Histology-MRI studies have demonstrated that MRI-derived grey matter atrophy reflects neurodegeneration (Filippi *et al*., 2012).

Grey matter atrophy is not uniform across the brain in MS and some regions are more susceptible to atrophy than others (Steenwijk *et al*., 2016; Preziosa *et al*., 2017). The limbic system, temporal cortex and deep grey matter show early atrophy in patients with relapse-onset MS (Audoin *et al*., 2010), whilst the cingulate cortex shows early atrophy in PPMS (Eshaghi *et al*., 2014). In our previous study using the same large cohort of MS patients (Eshaghi *et al*., 2017), we have found that the deep grey matter showed the fastest annual rate of tissue loss in relapsing-remitting MS and progressive MS, and that in the cortex the rate of atrophy accelerated in the temporal regions in secondary progressive MS. However, it is unknown whether there is a consistent and identifiable order in which atrophy progresses affecting different regions over time. A key question is whether there is an association between the sequential development of atrophy and disability accumulation.

One approach to investigate the sequence of atrophy progression is to employ a probabilistic data-driven method, such as an event-based model (EBM), which, as the name implies, identifies the sequence of events at which a biomarker becomes abnormal, using cross-sectional or longitudinal observations (Fonteijn *et al*., 2012; Young *et al*., 2014). The EBM is an established method. It has given new insights on the progression of Alzheimer’s in which the hippocampal atrophy is seen before the whole brain atrophy. Similarly in Huntington’s disease, the EBM has successfully predicted earlier atrophy in the basal ganglia than other regions (Fonteijn *et al*., 2012; Young *et al*., 2014).

In this study, we used the EBM to investigate the progression of brain atrophy as a sequence of “events” at which grey matter regions become atrophic in all phenotypes of MS. This method is data-driven; it does not rely on *a priori* thresholds to define when the volume of a region ceases to be normal and becomes atrophic, but calculates the probability of atrophy based on data-derived model distributions of normal and atrophic regional volumes. Moreover, the EBM constructs a subject staging system: it assigns each subject to a stage that reflects how far through the sequence of regions that subject shows lower than normal volumes - the higher the stage, the greater the number of atrophic regions.

In this study, we built on the evidence that neurodegeneration in MS does not affect all the grey matter regions equally (Haider *et al*., 2016; Eshaghi *et al*., 2017) and that brain regions become atrophic in a non-random manner (Rocca *et al*., 2010). We hypothesised that: (i) there is a sequence in which grey matter regions become atrophic; (ii) this sequence differs between relapse‐ and progressive-onset phenotypes; and (iii) the EBM stage increases with disease duration and disability worsening.

## Methods

### Participants

This was a retrospective study of 1,424 participants, studied between 1996-2016 in 7 European centres, which were part of the Magnetic Resonance in MS (MAGNIMS) Collaboration (www.magnims.eu). The same participants were previously used to investigate the spatiotemporal pattern of grey matter atrophy in MS (Eshaghi *et al*., 2017). They were healthy controls (HCs), patients with clinically-isolated syndrome (CIS), relapsing-remitting (RR) MS, secondary-progressive (SP) MS and primary-progressive (PP) MS. Eligibility criteria included: (1) A diagnosis of CIS or MS according to 2010 McDonald Criteria (Polman *et al*., 2011); (2) Healthy controls without history of neurological or psychiatric disorders; (3) The presence of at least 2 sequential MRI scans, acquired with identical protocol, including T1-weighted MRI and T2-weigthed/Fluid Attenuated Inversion Recovery (FLAIR) sequences; (4) A minimum interval of 6 months between MRI scans. We requested that the Expanded Disability Status Scale (EDSS) scored at clinical follow-ups on the eligible patients was made available (Kurtzke, 1983).

An additional group of age-matched healthy controls (N=29) was also obtained from the Parkinson’s Progression Markers Initiative (PPMI) database (www.ppmi-info.org/data) to match healthy control’s age to that of patients.

MRI scans were acquired under written consent obtained from each participant independently in each centre. The final protocol for this study was reviewed and approved by the European MAGNIMS collaboration for the analysis of pseudo-anonymised scans.

### MRI data and analysis

We collected 3D T1-weighted scans, in addition to T2/FLAIR MRI, from all centres except one. Details of the 13 different MRI protocols are shown in **Supplementary Table 1**.

The aim of the image analysis was to extract the volume of brain regions according to the Desikan-Killiany-Tourville protocol (Klein and Tourville, 2012) [as explained in detail elsewhere (Eshaghi *et al*., 2017)]. Briefly, the main steps were as follows: After an N4-bias field correction (part of ANTs software, version 4.1.9), which adjusted for the inhomogeneous intensity of the T1-weighted scans (Tustison *et al*., 2010), we performed T1 lesions filling (Battaglini *et al*., 2012) to improve the accuracy of the segmentation. We then created an unbiased, within-subject template, and linearly transformed all the subject-specific T1 scans to this symmetric space, using Freesurfer version 5.3 (Reuter *et al*., 2010; Reuter and Fischl, 2011; Reuter *et al*., 2012). In the symmetric space, we segmented the T1 scans in grey matter, white matter and CSF using the geodesic information flows or GIF software (part of NiftySeg, http://cmictig.cs.ucl.ac.uk/niftyweb/) (Cardoso *et al*., 2015). Finally, we calculated the regional volumes in the cortex and deep grey matter (the volume of the bilateral regions were averaged between the left and right hemisphere), the brainstem, white matter, cerebellum and lateral ventricles, according to the Desikan-Killiany-Tourville protocol (http://braincolor.mindboggle.info/index.html)(Klein and Tourville, 2012).

### The event-based model (EBM)

We used the EBM, as described previously in (Fonteijn *et al*., 2011, 2012; Young *et al*., 2014), to estimate the most likely sequence in which selected regions become atrophic over time (see below details on region selection). We also repeated the same analysis using all brain regions to test the dependence of our findings on the region selection.

The EBM assumes that a population of patients represents the whole trajectory of disease progression (Fonteijn *et al*., 2011) and reconciles cross-sectional or short-term longitudinal data into a picture of the whole disease course. We therefore created separate EBMs for: (1) relapse-onset patients (CIS, RRMS, SPMS); (2) progressive-onset (PPMS) patients; and (3) to develop a unique staging system for all patients we merged all clinical phenotypes into a single cohort. We used the sequence estimated by this EBM to stage patients by assigning them the most probable stage along the sequence.

The main steps of the EBM include (**Figure 1**): (1) model input, which consists of adjustment of regional volumes for effects of nuisance variables and selection of regions; (2) model fitting; and (3) a cross-validation. For the last step, we used a novel cross-validation method, used here within the EBM for the first time, whilst steps 1 and 2 have not changed since the original EBM implementation (Fonteijn *et al*., 2011; Young *et al*., 2014). Model input used all MS patients; model fitting and cross-validation were repeated three times using (1) relapse-onset and CIS patients together, (2) PPMS, and (3) a merged cohort of patients.

**Figure 1.**
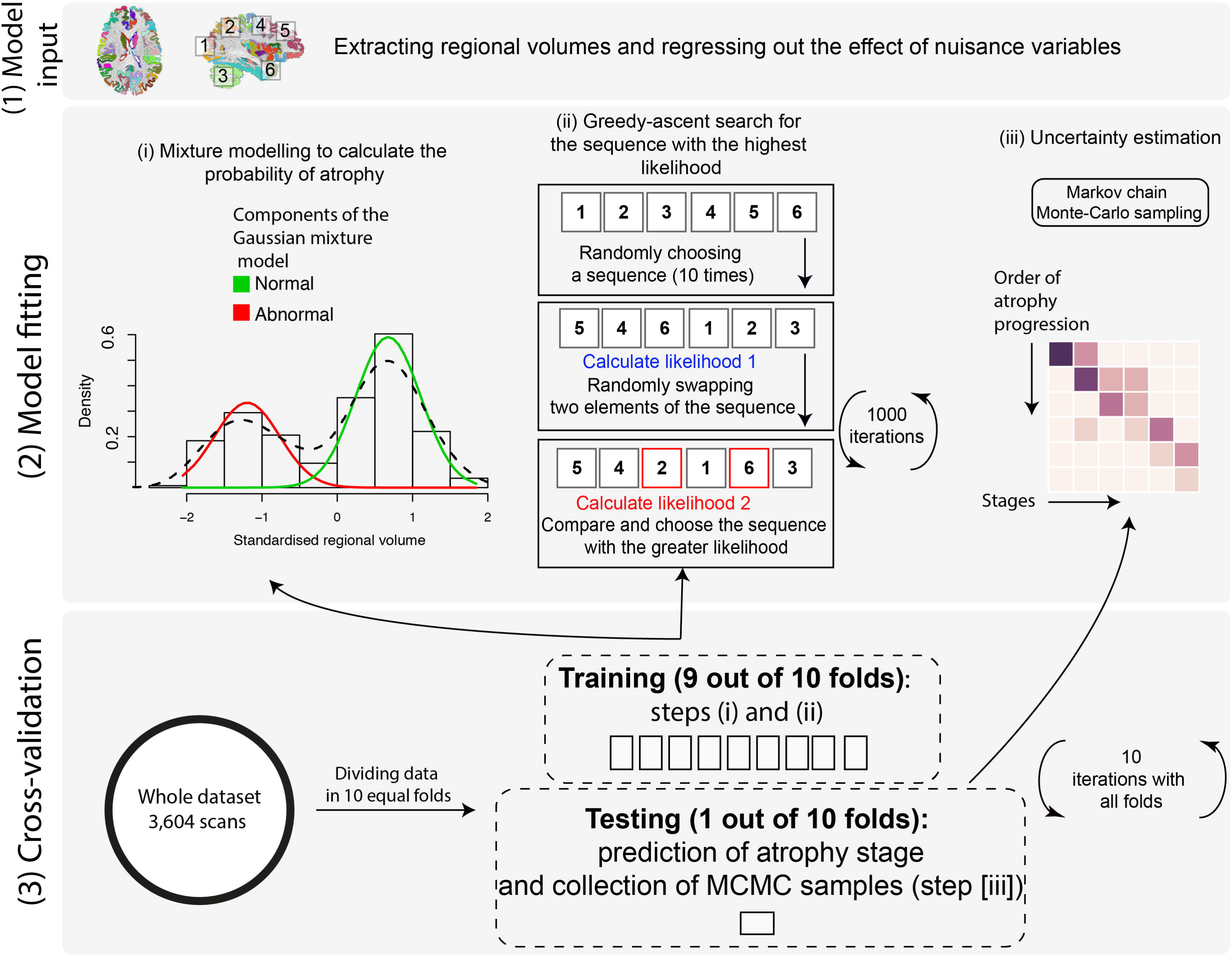
EBM steps to estimate the most likely sequence of atrophy progression. The 3 steps are: (1) calculating the best-fit probability distributions for normal and atrophic brain regions; (2) searching for the most likely sequence; and (3) quantifying the uncertainty with cross validation. (1) shows the distribution of the volume in an example region in healthy controls and patients. (2) shows the steps for greedy ascent search. (3) a matrix showing a sequence of atrophy progression on the *y*-axis, and the position in the sequence of each region ranging from 1 to the total number of regions on the *x*-axis. The intensity of each matrix entry corresponds to the proportion of Markov Chain Monte Carlo samples of the posterior distribution where a certain region of *y*-axis appears at the respective stage of *x*-axis.

### Model input

We adjusted the regional volumes for the total intracranial volume, age at study entry, gender, scanner magnetic field and MRI protocol. Since some centres provided data from more than one MRI protocol we adjusted for MRI protocol and magnetic field instead of only a “centre” variable (see Supplementary Table 1). We constructed a regression model for each region separately, entering the volume as the dependent variable and the remaining variables as predictors. We extracted the amount of each regional volume that remained unexplained in the regression (residual of the fit). Subsequently, we selected the regions whose adjusted volumes at the study entry showed a significant difference between all MS patients and healthy controls, with a Bonferroni corrected *p*<0.01 (non-corrected *p* <0.0001). We used these regions in the subsequent analyses. We repeated the analysis using all the segmented regions of the Desikan-Killiany-Tourville atlas for the following reasons: 1) To test whether the sequence in which brain regions become atrophic was not influenced by restricting the analysis only to the regions that showed a lower volume in patients than controls; 2) to detect potential subtle early changes that might have not survived multiple-comparison correction.

### Model fitting

The EBM considers an “event” to have occurred when a biomarker, here regional volume, has abnormal value (“atrophy”) in comparison with the expected values measured in healthy controls. The model then estimates the sequence *S* = *S(1), S(2), …, S(l)* in which regions become atrophic, where *S(1)* is the first region, and *S(l)* is the last to become atrophic. The model assumes that all patients go through the same sequence as they progress. The estimation procedure first fits a mixture of two Gaussians to regional volumes, with one of the components fixed to be identical to the healthy distribution; the other component provides the model for the “abnormal” distribution. This provides probabilistic models for normal and abnormal volumes from which we can calculate the likelihood of atrophy *P*(*x*_*ij*_|*E*_*i*_) for the region *i* of the scan *j*, i.e. the probability density function (PDF) estimated at *x*_*ij*_ from the abnormal component of the mixture-model. The likelihood that region *i* has no atrophy, or *P*(*x*_*ij*_|¬*E*_*i*_), is the PDF of the normal component of the mixture-model estimated at *x*_*ij*_ (see **Figure 1**, section 2[i]).

To search for the most likely sequence, we used a greedy ascent search (Fonteijn *et al*., 2012; Young *et al*., 2014) which started at 10 different random sequences and iterated by randomly flipping sequences for 1000 times. The final sequence was selected when 10 different initial sequences converged to a similar likelihood after 1000 iterations. Within each iteration new (flipped) sequences (Figure 1, section 2[ii]) were accepted only if they increased the likelihood, which is defined as

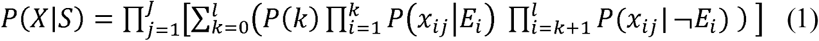

where *X* is the data matrix, *S* is the sequence of atrophy events, *J* is the number of scans, *l* is the number of regions, and *P(k)* is the prior probability of being at stage *k*, which means *E*_*i*_, …, *E*_*k*_ have occurred, and *E*_*k*+*1*_, …, *E*_*l*_ have not occurred. We used a uniform distribution for prior probabilities, which assumes equal prior-probability for all possible stages; all sequences are equally likely *a-priori*.

### Cross-validation of atrophy sequence

After estimating the most likely sequence, the uncertainty in the position of each region in the sequence was estimated using cross-validation and Markov Chain Monte-Carlo (MCMC). We divided the dataset (including baseline and follow-up visits) into ten equally-sized folds (cross-validation folds) and repeated the sequence estimation ten times. During each iteration, we used nine-folds to fit the mixture-models (as explained above) and estimated the most-likely sequence. We kept one fold out as the test fold to assign the EBM stages (explained below). Within each iteration, we used MCMC to sample from the posterior distribution on the sequence given the nine-fold training data, as in (Fonteijn *et al*., 2012; Young *et al*., 2014). We then aggregated MCMC samples from the 10 iterations of cross-validation (10,000 samples from each fold) to calculate uncertainty across cross-validation folds. Finally, we used these 100,000 sampled sequences to plot the positional variance diagram (as in (Fonteijn *et al*., 2012; Young *et al*., 2014)), which shows on the y-axis the sequence with the highest likelihood, and the *x*-axis enumerates the number of sequence positions (or EBM stages). The intensities of the matrix entries correspond to the proportion of MCMC samples in which the corresponding region (*y*-axis) appears at the respective stage (*x*-axis). Therefore, if there were no uncertainty, i.e. all MCMC samples in all folds find the same sequence, the matrix would be black on the diagonal and white everywhere else; non-white off-diagonal and non-black diagonal elements indicate uncertainty in the position of the corresponding region in the sequence.

### Staging individual subjects and clinical associations

We used the most likely sequence of atrophy progression from a unique EBM created from a merged patient cohort to obtain the EBM stage for each MRI scan *j*, which is the stage *k* that maximises 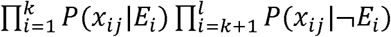. This assigned each subject an EBM stage between 1 and the number of regions, *l*, at each visit (see **Figure 1**).

We used a nested linear mixed-effects regression model to explore the associations between changes of EBM stage over time with disease duration, where disease duration was nested in subjects (random-effects), and disease duration was also a fixed-effects variable.

For those clinical phenotypes that showed a significant change in the EBM stage over time (RRMS, SPMS and PPMS), we investigated whether longitudinal EDSS changes could be predicted by EBM changes independent of disease duration. We divided the changes in EDSS and EBM by number of years from the study entry, and performed a linear regression analysis where annualised EDSS change was the outcome variable. Annualised EBM change and disease duration at the study entry were the predictor variables. Since both the EBM stage and EDSS are ordinal variables, we used ordinal regression analyses to confirm the results of the linear regressions but presented the results of linear models (as they did not materially differ).

## Results

### Subject characteristics

Imaging data from 1,424 subjects were analysed; three subjects’ scans were excluded because of motion artefacts and four because of poor registration due to missing MRI header information. Therefore, data from 1,417 subjects were included in the final modelling: 1,214 patients (253 CIS, 708 RRMS, 128 SPMS, and 125 PPMS), and 203 healthy controls (HCs). The average (±standard deviation) length of follow-up for patients was 2.43 years (±1.97) and for HCs was 1.83 years (±1.77). In total, we analysed 3,604 T1-weighted MRI scans (mean number of scans per patient was 2.54 [SD=1.04]).

**Table 1.** Baseline characteristics of participants.

| Group | Healthy control | CIS | Relapse onset | PPMS |
| --- | --- | --- | --- | --- |
| subgroups |  |  | RRMS & SPMS |  |
| Total number<br>(number of females) | 203 (112) | 253 (171) | 836 (548) | 125 (55) |
| Age (± SD<sup>1</sup>) | 38.7 ± 10.5 | 33±8 | 39.7±9.8 | 48.5 ± 10.1 |
| Disease duration (± SD) | — | 0.4±1.4 | 8.06±8.03 | 6.8 ± 5.9 |
| Median EDSS<sup>2</sup><br>(range) | — | 1 (0-4.5) | 2 (0-9) | 5 (2-8) |
| Percent (number)<br>receiving disease<br>modifying drugs | — | 20% (52) | 397 (47%) | 6% (8) |

### Sequence of atrophy progression

At baseline, 24 regions showed a smaller volume in MS than HCs with a Bonferroni corrected *p*<0.01. They included the deep grey matter regions and the posterior cortices (including the precuneus and the posterior cingulate cortex), several regions in the temporal lobe, the precentral cortex, and the brainstem (see **Figure 2** for the full list).

**Figure 2.**
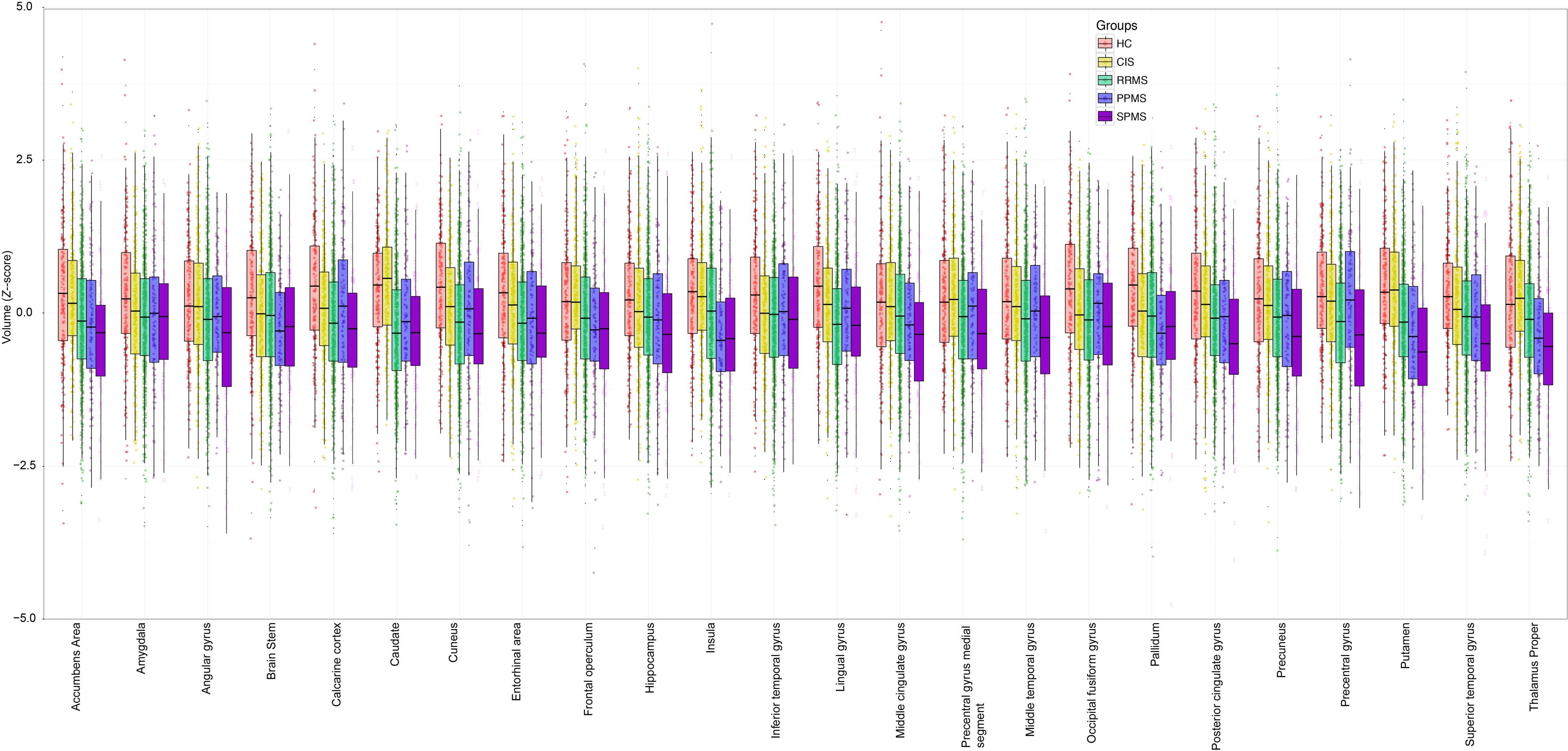
Comparisons of regional volumes between groups. Box plots at *y*-axis show *z*-scores of the corresponding region shown at *x*-axis. Lower and upper hinges of each boxplot correspond to 25^th^ and 75^th^ percentiles of data. We arbitrarily selected 24 regions (shown in A) according to the *t*-value of the comparison between all patients with MS and healthy controls at baseline visit.

When we estimated the sequence in which these 24 regions become atrophic in patients with relapse-onset MS (RRMS and SPMS) and CIS, the first regions were the posterior cingulate cortex and the precuneus, followed by the middle cingulate cortex, brainstem, and thalamus (**Figure 3A & D**); the last regions to become atrophic were the pallidum, and medial precentral gyrus.

**Figure 3.**
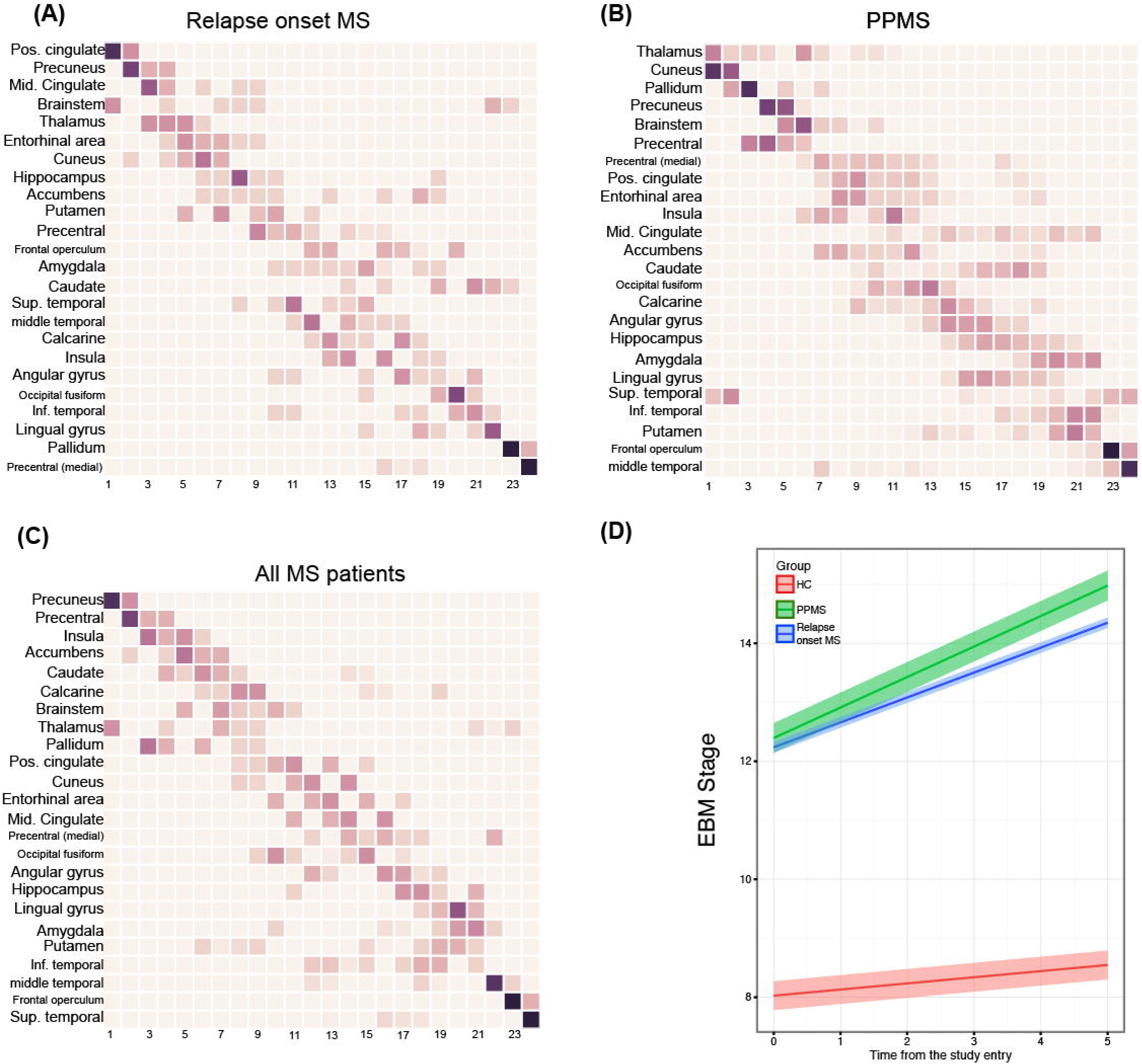
Sequences of atrophy progression and patient staging. The positional variance diagrams for (A) relapse-onset MS, (B) PPMS and (C) all patients together show the most likely sequences of atrophy and their associated uncertainty. In (A), (B), and (C) the *y*-axis shows the most likely sequence of atrophy progression, and the *x*-axis shows the sequence position ranging from 1 to the total number of regions. The intensity of each rectangle corresponds to the proportion of Markov Chain Monte Carlo samples of the posterior distribution where a certain region of *y*-axis appears at the respective stage of *x*-axis. (D) shows the evolution of EBM stage (or atrophy progression staging) over time in CIS and relapse-onset MS together and PPMS. Each line in (D) is the prediction of mixed-effects model whose ribbon shows standard error of the prediction.

In patients with PPMS, among the 24 selected regions, the first ones to show atrophy were the thalamus, cuneus and precuneus, and pallidum, followed by the brainstem, precentral gyrus, and posterior cingulate cortex (**Figure 3B & D**); the last regions to become atrophic were the frontal operculum and middle temporal gyrus.

When the EBM was used to estimate the sequence of atrophy progression of the selected 24 regions in all patients together, additional regions were detected as showing early atrophy, such as the insula, accumbens and caudate (**Figure 3C**). The likelihood of the 10 randomly chosen sequences (log-likelihood range: -149000 to -117000) converged to a similar range (log-likelihood range: -1000000 to -99000) after 1000 iteration (**Supplementary Figure 1**). For other EBMs, the likelihoods converged to a similar range (results are not shown).

When all the remaining regions were included additional regions were identified. In PPMS they were the transverse temporal gyrus, cerebral white matter, post-central gyrus and middle frontal gyrus (see **Figure 4, Supplementary Figures 2** and **3**). In relapse-onset group these regions were the superior frontal gyrus, inferior frontal gyrus, and middle frontal gyrus.

When we qualitatively compared CIS and relapse-onset MS patients with PPMS, across all regions, the cerebellum, caudate and putamen showed a differential pattern of atrophy, with early atrophy in patients with relapse-onset disease and late atrophy in PPMS (see **Figure 4**).

**Figure 4.**
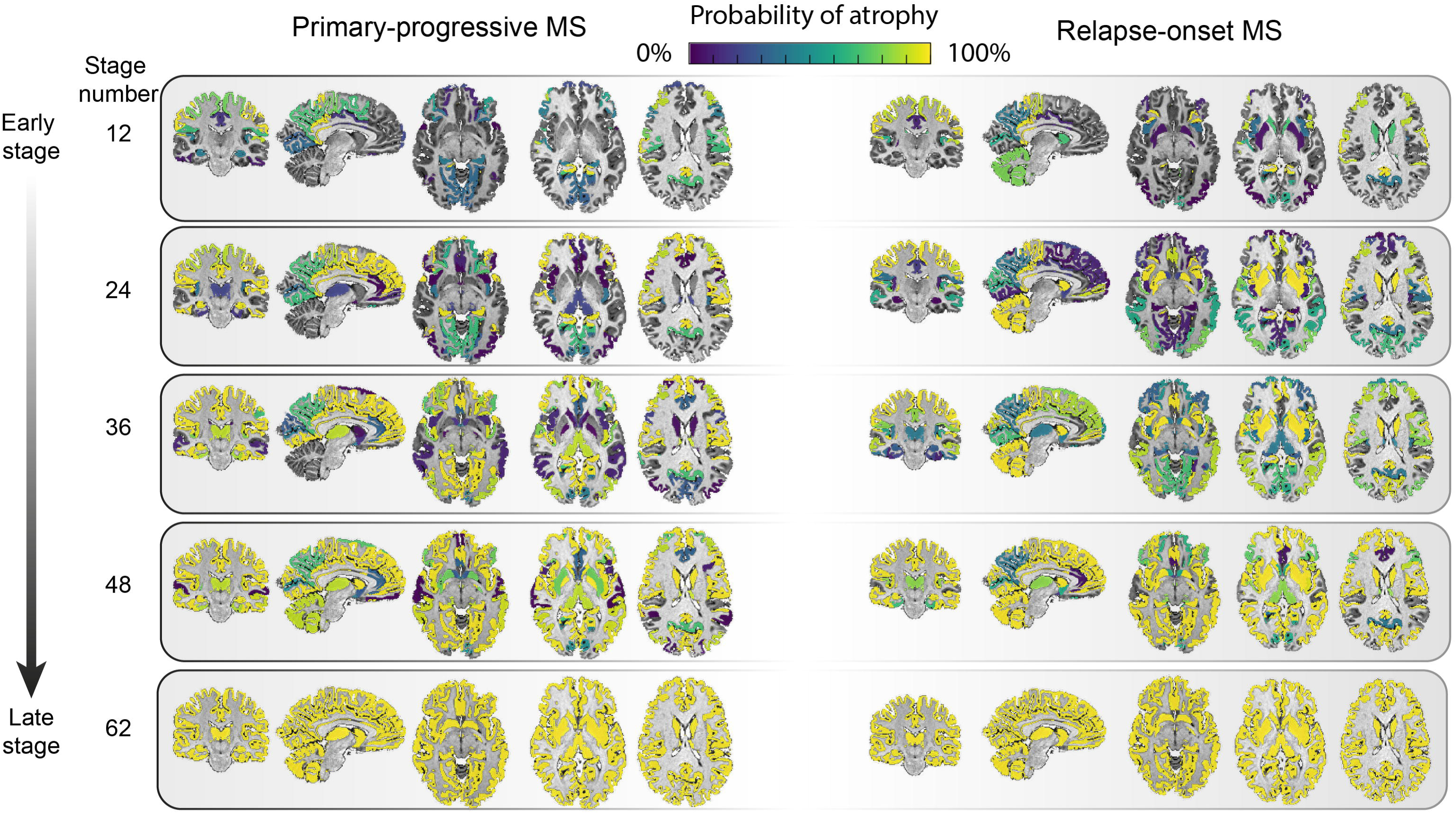
Regional atrophy and its sequence of progression in all grey matter regions plus brainstem in relapse-onset disease and PPMS. The probability of atrophy in each region was calculated from the positional variance diagrams and colour coded, so that brighter colour corresponded to higher probability of seeing atrophy in the corresponding EBM stage.

### EBM staging of individual subjects

Patients with relapse-onset MS and PPMS had significantly higher EBM stages at baseline than HCs (average intercept [±standard error] EBM stage for HCs=8.02±0.59, relapse-onset=12.39±0.66, PPMS=12.22±0.35, *p*<0.05). When looking at each clinical phenotype, patients with SPMS had the highest EBM stage at the study entry than other clinical phenotypes (average intercept ± standard error=14.73±0.93, all *p*-values<0.001), followed by RRMS (12.60±0.67), PPMS (12.22±0.35), CIS (8.12±0.76), and HCs (8.02±0.59) respectively. The annual rate of change (or slope) of the EBM stage over time was significant (null hypothesis=zero slope) for SPMS (average slope± standard-error:1.02±0.41), PPMS (0.52±0.34), and RRMS (0.37±0.26), but not for CIS (0.19±0.33) or HCs (0.10±0.24). The rate of change, although nominally higher in SPMS, was not significantly different between clinical phenotypes.

### Associations between EBM staging and disease duration and disability accumulation

There was a significant association between the rate of increase in the EBM stages and disease duration in all patients with MS (*β*=0.21, standard error [SE]=0.03, *p*<0.001) using all available time points. This means that for every increase of one EBM stage, disease duration increased by 4.76 years.

At the baseline visit, there was no significant association between the EBM stage and EDSS in any clinical phenotype (CIS, RRMS, SPMS, or PPMS). Over time there was a significant increase in EDSS in both relapse-onset/CIS and PPMS patients (increase of 0.07 and 0.2 per year, respectively, *p*<0.01). There was a significant association (independent of disease duration) between annualised EBM stage and annualised EDSS changes in RRMS (beta =0.03, p <0.0001) but not in SPMS and PPMS. This means that assuming a linear relationship between EDSS and EBM stage, for every unit increase in the annual rate of EBM stage there is 0.03 increase in the annual rate of EDSS worsening. There was no significant association between the rate of change in EBM stage and lesion load over time.

## Discussion

In this study, we used a data-driven method to determine the most likely sequence in which brain regions become atrophic in MS. This sequence is consistent in key regions across MS phenotypes: the posterior cingulate cortex, the precuneus, and the thalamus were among the earliest regions to become atrophic in both relapse-onset phenotypes and PPMS. The EBM predicted a different pattern in the cerebellum, caudate, and putamen, which was an early atrophy in relapse-onset but a later atrophy in PPMS. The EBM staging system was applied to individual patients and the rate of increase in the EBM stage was associated with the disease duration in all MS phenotypes and with the EDSS in patients with RRMS independent of disease duration. These results provide insights into the mechanisms of disease worsening in MS.

The order of atrophy progression in the EBM for most regions was similar between PPMS and CIS/relapse-onset MS. This may support the evidence from histological studies that the pathological processes are regionally consistent between early RRMS and progressive MS (SPMS and PPMS) (Mahad *et al*., 2015). Our results showed that areas with an early atrophy according to the EBM were the posterior cingulate cortex, precuneus, thalamus and brainstem in both groups, thereby extending the results of previous studies, which have limited their investigation to specific MS subtypes (Gilmore *et al*., 2009; Audoin *et al*., 2010; Calabrese, Reynolds, *et al*., 2015; Steenwijk *et al*., 2016). When all patients were included together, the insula, the accumbens and the caudate were predicted as becoming atrophic early on.

The cingulate cortex and insula are interconnected with other regions, and vulnerable to secondary damage via white matter tracts (Bodini *et al*., 2016). Moreover, these structures are in cortical invaginations, which could expose them to meningeal inflammation, cortical demyelination, and neurodegeneration (Gilmore *et al*., 2009; Howell *et al*., 2011; Haider *et al*., 2016). The cingulate cortex and precuneus are part of a network of active regions during rest (the default mode network) (Raichle, 2015). These regions are interconnected with other areas, have the highest energy consumption in the brain, and are affected in MS and other neurodegenerative disorders (Bonavita *et al*., 2011; Raichle, 2015). In MS, neurons with demyelinated axons consume more energy to adapt to demyelination, which creates a microenvironment similar to that of hypoxia (“virtual hypoxia”) (Trapp and Stys, 2009). Neurons that survive in a state of persistent virtual hypoxia are more vulnerable to degeneration (Zhang and Raichle, 2010), and this may explain higher vulnerability of the cingulate and precuneus cortex to atrophy.

Other regions that showed early atrophy were the thalamus and the brainstem in both relapse-onset MS and PPMS. In our previous study, we found that the deep grey matter showed the fastest rate of atrophy over time, whilst brainstem had the highest atrophy (the lowest volume) at study entry, but its atrophy progressed at a slower rate than that occurring in other regions (Eshaghi *et al*., 2017). This may suggest that during early stages of MS, the rate of atrophy in the brainstem is higher than later stages, whilst the rate of atrophy in the thalamus remains high throughout the disease course. The brainstem is in close contact with the spinal cord, whose atrophy is seen from early stages of MS independent of the cortex or deep grey matter (Biberacher *et al*., 2015; Ruggieri *et al*., 2015).

Several mechanisms may underlie neurodegeneration in the deep grey matter, including mitochondrial failure, iron deposition, retrograde degeneration through white mater lesions, and meningeal inflammation (for structures closer to CSF) (Calabrese, Magliozzi, *et al*., 2015; Bodini *et al*., 2016; Haider *et al*., 2016; Pardini *et al*., 2016). Network overload and collapse, similar to the cingulate and precuneus cortex, could also explain preferential atrophy of the deep grey matter in MS (Minagar *et al*., 2013).

There were a few regions showing differential pattern of atrophy between relapse‐ and progressive-onset phenotypes. The cerebellum, caudate and putamen were predicted to have early atrophy in relapse-onset disease and late atrophy in PPMS. In the cerebellum this different behaviour can be explained by a more inflammatory phenotype of patients with relapse-onset MS. In patients with MS, more than any other brain region, demyelination is seen in the cerebellar grey matter, which is 5 times more than the white matter demyelination (Gilmore *et al*., 2009). This may be a consequence of overlying meningeal inflammation in the deep folia, which accommodate a static inflammatory milieu (such as cytokines, and immunoglobulins) (Kutzelnigg *et al*., 2007; Howell *et al*., 2011). Therefore, in the cerebellum overlying inflammation may play a role and amplify other pathological mechanisms, such as retrograde neurodegeneration secondary to white matter lesions (Kutzelnigg *et al*., 2007; Gilmore *et al*., 2009; Howell *et al*., 2011). Thus, the cerebellum could be susceptible to inflammatory damage from CSF. Previous studies have reported in relapse-onset MS, but not PPMS, tertiary lymphatic follicles in cortical invaginations, which may suggest a more inflammatory CSF milieu than PPMS (Kutzelnigg *et al*., 2007; Choi *et al*., 2012). This could explain earlier atrophy of the cerebellar grey matter in people with relapse onset disease, whilst in PPMS neurodegeneration in a less inflammatory CSF milieu might cause a gradual progression of atrophy (Choi *et al*., 2012; Mahad *et al*., 2015). However, this is speculative and it remains unclear whether meningeal inflammation has a causative effect on demyelination and neurodegeneration.

The caudate and putamen, which are histologically similar, constitute a structure that is known as the neostriatum. A previous histopathological study has shown that the highest extent of demyelination and lesions in the deep grey matter can be seen in the caudate even in early MS, although the pattern was not different between MS phenotypes (Haider *et al*., 2014). Moreover, the neostriatum receives major inputs to the deep grey matter from the motor cortex (mostly to putamen) and the association cortices (mostly to the caudate). Therefore, we could speculate that retrograde neurodegeneration secondary to a higher lesion load in relapse-onset disease (compared to PPMS) may perform as an additive factor on demyelination to explain the higher vulnerability of these structures.

We extended our analysis from regions that showed significant atrophy at baseline to the all segmented regions to test the dependence of our findings to region selection. Another reason was to explore early, but subtle, changes in brain regions, which might have been missed by just looking at a snapshot at the study entry to choose specific regions based on stringent multiple-comparison correction. For example, a brain region may show mild volume loss earlier than another region with a greater (but later) volume loss through the course of MS. Whole brain EBM analysis predicted an early involvement of the posterior cortices (posterior cingulate and precuneus) along with the brain stem. New additional regions in the whole brain EBM were also identified including: the superior, middle, and inferior frontal gyri in relapse-onset phenotypes, and the transverse temporal gyrus, white matter, and post-central gyrus in PPMS. These findings suggest that the changes in these structures may happen early, but with a lower intensity than other regions that were only reported in the EBM with 24 regions.

The EBM has potential for clinical use as it does not rely on time, and can be applied to individual (cross-sectional) MRI scans. To have a unique staging system across clinical phenotypes, we therefore created a separate EBM from a merged MS cohort. We showed that patients with SPMS had the highest EBM stage ‐or the highest number of atrophic regions-at the study entry. This, in line with previous studies, suggests that SPMS has more advanced neurodegeneration across MS phenotypes (Ceccarelli *et al*., 2008). When we performed the EBM staging using follow-up scans of patients and healthy controls, we found a significant increase in EBM stages in all MS phenotypes, but not in CIS or HCs (although the baseline EBM stage was nominally higher in CIS than HCs). The clinical relevance of the EBM was confirmed by a significant association between stages and EDSS after adjusting for disease duration in RRMS. Therefore, the sequential pattern of atrophy may explain disease worsening in RRMS. We did not find the same association between the changes in EBM stages and EDSS. However, patients with SPMS had the highest EBM stages at the study entry, and the highest nominal rate of increase in the EBM stage (although not significantly so in comparison with RRMS or PPMS). This suggests that sequential pattern of atrophy may explain mechanisms of disease progression.

Although this is a retrospective and multi-centre study, we have adjusted for the effects of centre; as reported before on this dataset (Eshaghi *et al*., 2017), the effect of MS phenotypes on regional measures was higher than that from the centre. A possible limitation is that EBM assumes that all brain regions eventually become abnormal (all regions show atrophy at the last stage). Therefore, an implicit assumption is that patients with relapse-onset disease (CIS, RRMS, and SPMS) or those with PPMS, represent the whole continuum of progression when analysed separately; future implementations of this model could remove this assumption. We used EDSS as the clinical outcome, but both EDSS and EBM provide measures that are ordinal, and may not have a uniform interpretation. Therefore, the coefficients of associations should be interpreted relatively (e.g., to compare clinical groups) rather than absolutely.

We showed that the sequence of atrophy progression in relapse-onset disease and PPMS are similar in many key regions, whilst the cerebellum, caudate and putamen show an earlier atrophy in relapse-onset MS, perhaps due to a more inflammatory milieu. The sequence of atrophy progression can be used to score automatically patients during their path of atrophy progression, and has potential clinical utility because for staging of patients.

## Acknowledgements

Authors acknowledge The National Institute for Health Research (NIHR) Biomedical Research Centre (BRC) at University College London Hospitals (UCLH) for their support. A Eshaghi receives McDonald Fellowship from Multiple Sclerosis International Federation (MSIF, http://www.msif.org). He has received a fellowship from the MAGNIMS collaboration (http://www.magnims.eu). PPMI – a public-private partnership–is funded by the Michael J. Fox Foundation for Parkinson’s Research and funding partners (see the full list at www.ppmi-info.org/fundingpartners). This project has received funding from the European Union’s Horizon 2020 research and innovation programme under grant agreement No 666992.

## Figure titles

Figure 1. Estimating the most likely sequence of atrophy progression.

Figure 2. Regional volumes of the chosen regions across groups at baseline.

Figure 3. Most likely sequences of atrophy progression and patient staging.

Figure 4. Regional atrophy and its sequence of progression.

## Supplementary Legends for Figures

**Supplementary Figure 1:**
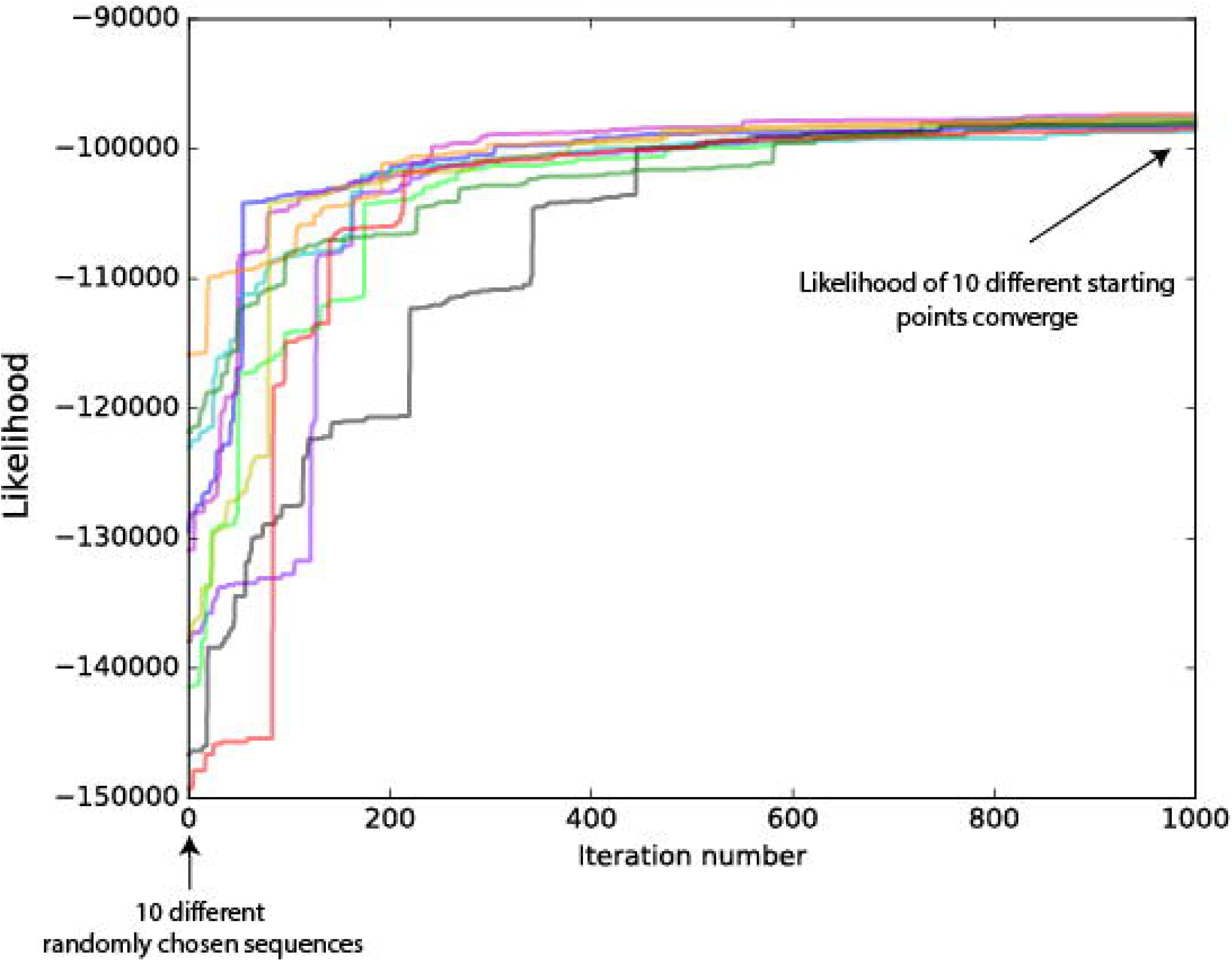
Greedy ascent search and convergence of likelihoods. The most likely sequence of atrophy progression in relation to 10 randomly chosen initial sequences. The *y*-axis shows the data likelihood (calculated from Equation 1). The *x*-axis shows the number of iterations at which two events are randomly swapped in search for a higher sequence likelihood. This procedure was repeated during each cross-validation (10 times).

**Supplementary Figure 2.**
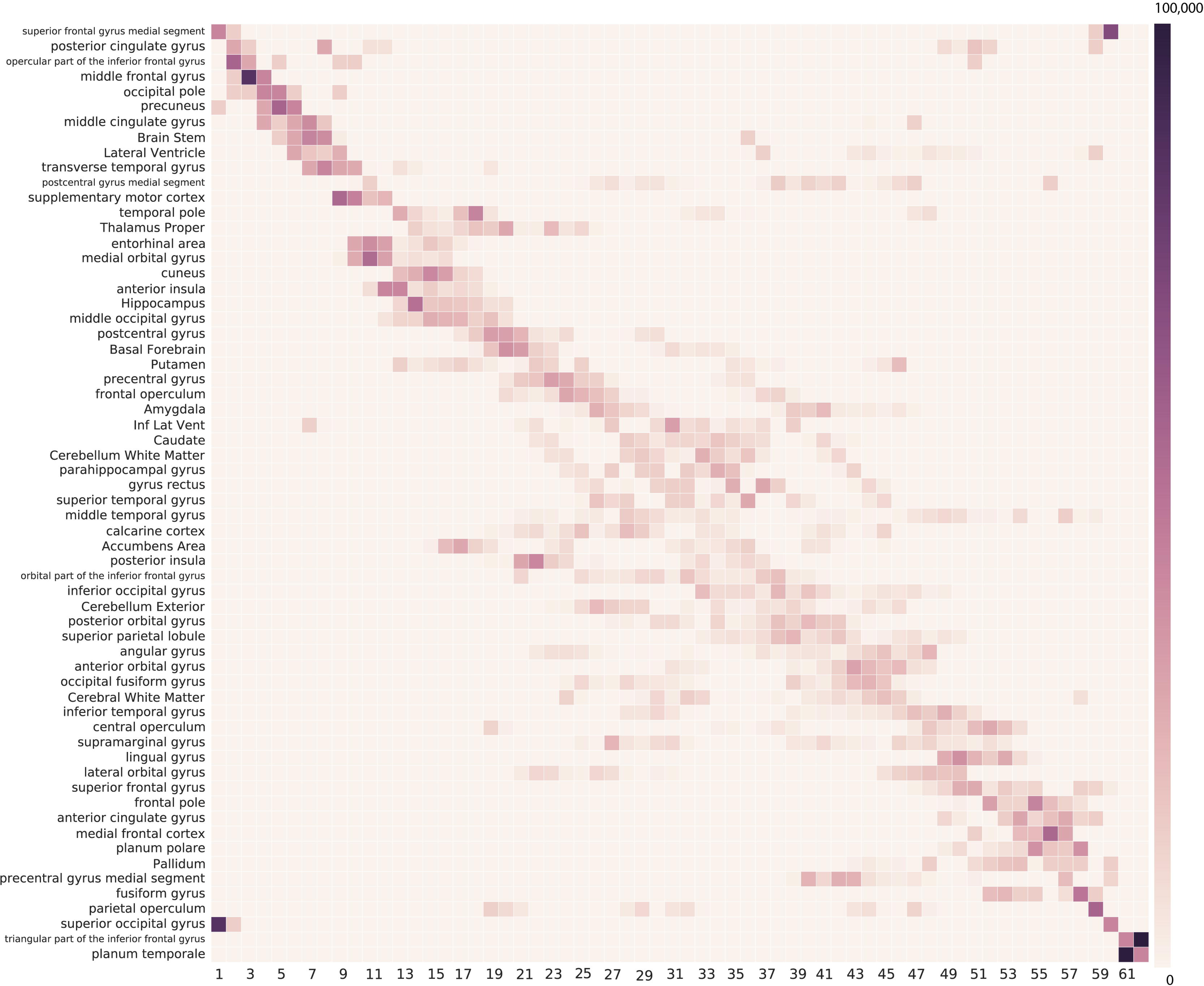
Event-based model applied to all regions in patients with CIS/relapse-onset MS.

**Supplementary Figure 3.**
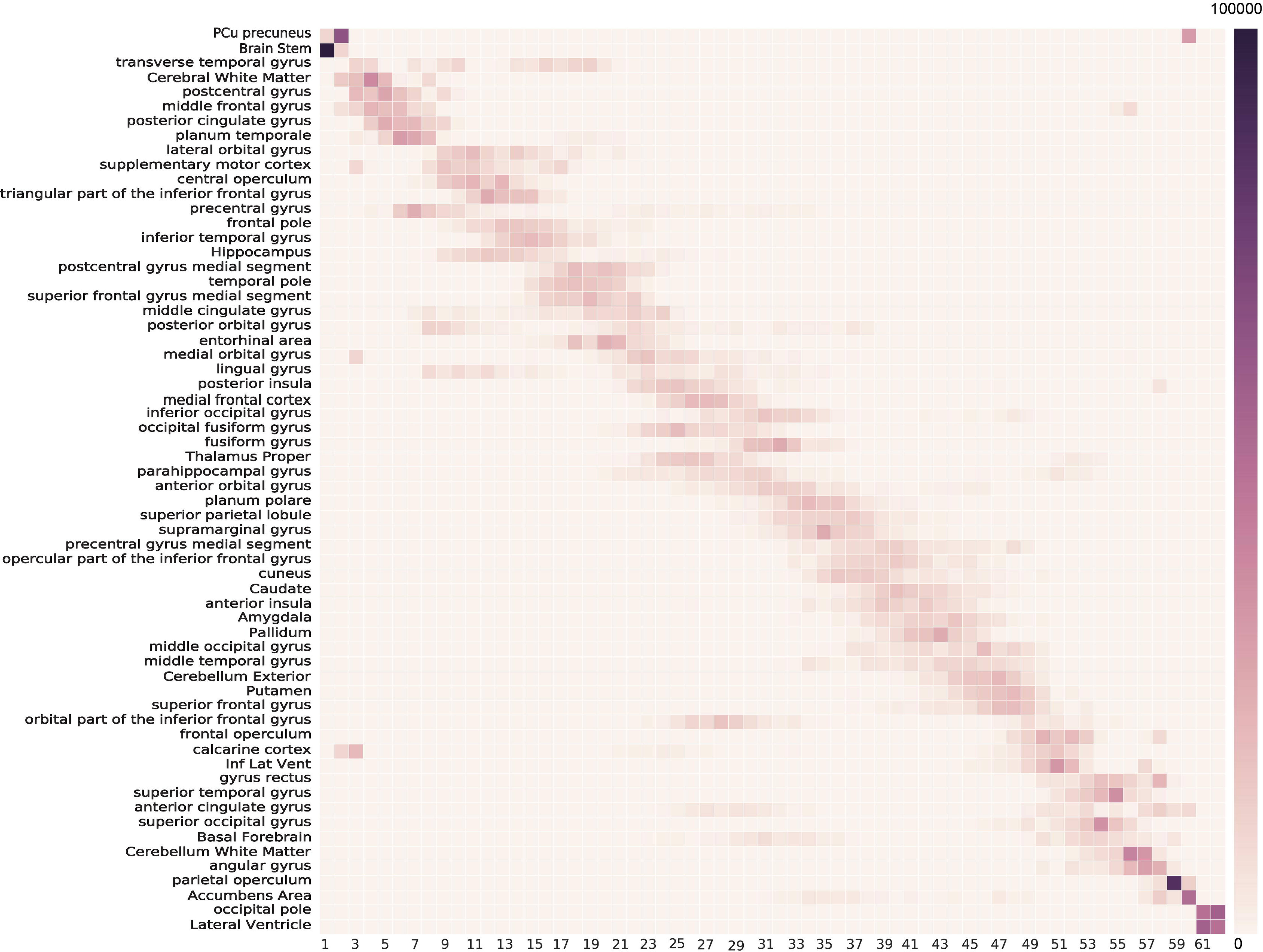
Event-based model applied to all regions in patients with PPMS.

## Supplementary figure titles

Supplemental Figure 1. Greedy ascent search.

Supplementary Figure 2. Positional variance diagram for CIS/relapse-onset MS based on all brain regions.

Supplementary Figure 3. Positional variance diagram for PPMS based on all brain regions.

**Supplementary Table 1.** MRI protocols for each participating centre.

|  | T1-weighted MRI |  |  |  |  |  |  | Sequence for lesion delineation |  |  |  |  |  |
| --- | --- | --- | --- | --- | --- | --- | --- | --- | --- | --- | --- | --- | --- |
|  | Magnetic field | Vendor | Voxel dimension | TR | TE | Matrix size | Slices | MR Sequence | TE | TR | TI | Matrix size | Slice thickness |
| **London** | 1.5T | General Electric Signa | 3D<br>(1.2x1.2x1.5 mm) | 13.3 ms | 4.2 ms | 256x256 | 124 | T2 | 80 ms | 1720 ms | – | 256x256 | 5 mm |
|  | 1.5T | General Electric Signa | 3D<br>(1.2x1.2x1.2) | 14.3 ms | 5.1 ms | 256x256 | 156 | PD-T2 | 17-102 ms | 2000 ms |  | 256x256 | 5 mm |
|  | 1.5T | General Electric Signa | 3D<br>(1.2x1.2x1.5) | 29 ms | 15 ms | 256x256 | 124 | PD-T2 | 17-102 ms | 2000 ms | – | 256x 256 | 5 mm |
|  | 3T | Philips Achieva | 3D<br>(1x1x1 mm) | 6.8 ms | 3.1 ms | 256x256 | 256 | PD-T2 | 19-85 ms | 3500 ms | – | 240x240 | 3 mm |
| **Milan** | 1.5T | Siemen Avanto | 3D<br>(1x1x1 mm) | 2000 ms | 3.93 ms | 256*224 | 208 | PD-T2 | 28-113 ms | 2560 ms | – | 256x256 | 2.5 mm |
|  | 3T | Philips Intera | 3D<br>(0.89x0.89x1 mm) | 25 ms | 4.6 ms | 256*256 | 220 | PD-T2 | 24-120 ms | 3350 ms | – | 256x256 | 3 mm |
| **Graz** | 3T | Siemens Tim Trio | 3D<br>(1x1x1 mm) | 1900 ms | 2.6 ms | 176*221 | 256 | FLAIR | 69 ms | 10000 ms | 2500 ms | 192x256 | 3 mm |
| **Barcelona** | 1.5T | Siemens Symphony | 3D<br>(1x1x1 mm) | 1980 ms | 3.1 ms | 256x256 | 176 | FLAIR | 95 ms | 8500 ms | 2440 ms | 192x256 | 3 mm |
|  | 3T | Siemens Tim Trio | 3D<br>(1x1x1.2 mm) | 2300 ms | 2.98 ms | 256x240 | 128 | FLAIR | 93 ms | 9000 ms | 2500 ms | 400x512 | 3 mm |
| **Amsterdam** | 1.5T | Siemens Vision | 3D<br>(1x1x1 mm) | 4000 ms | 20 ms | 180x256 | 256 | T2 | 20 ms | 4000 ms | 108 ms | 256x256 | 3 mm |
| **Rome** | 1.5T | Siemens Avanto | 3D<br>(1x1x1 mm) | 9000 ms | 89 ms | 192x256 | 160 | FLAIR | 89 ms | 9000 ms | 2500 ms | 192x256 | 3 mm |
| **Siena** | 1.5T | Philips Gyroscan | 2D<br>(0.97x0.97x3mm ) | 35 ms | 10 ms | 256x256 | 50 | FLAIR | 150 ms | 9000 ms | 2725 ms | 256x256 | 3mm |
| **PPMI** | 3T | Siemens Tim Trio | 3D<br>(1x1x1mm) | 2300 ms | 2.52 ms | 176x240 | 256 | – | – | – | – | – | – |

## Appendix

*MAGNIMS Steering Committee Members

Alex Rovira: MR Unit and Section of Neuroradiology, Department of Radiology, Multiple Sclerosis Centre of Catalonia (CEMCAT), Hospital Universitari Vall d'Hebron, Universitat Autònoma de Barcelona, Barcelona, Spain

Christian Enzinger: Department of Neurology, Medical University of Graz, Graz, Austria

Frederik Barkhof: Queen Square Multiple Sclerosis Centre, UCL Institute of Neurology, University College London, London, UK

Olga Ciccarelli: Queen Square Multiple Sclerosis Centre, UCL Institute of Neurology, University College London, London, UK

Massimo Filippi: Neuroimaging Research Unit, Institute of Experimental Neurology, Division of Neuroscience, San Raffaele Scientific Institute, Vita-Salute San Raffaele University, Milan, Italy

Nicola De Stefano: Department of Medicine, Surgery and Neuroscience, University of Siena, Siena, Italy

Ludwig Kappos: Department of Neurology, University Hospital, Kantonsspital, Basel, Switzerland

Jette Frederiksen: The MS Clinic, Department of Neurology, University of Copenhagen, Glostrup Hospital, Denmark

Jaqueline Palace: Centre for Functional Magnetic Resonance Imaging of the Brain, University of Oxford, UK Maria A Rocca: Neuroimaging Research Unit, Institute of Experimental Neurology, Division of Neuroscience, San Raffaele Scientific Institute, Vita-Salute San Raffaele University, Milan, Italy

Jaume Sastre-Garriga: Department of Neurology/Neuroimmunology, Multiple Sclerosis Centre of Catalonia (CEMCAT), Hospital Universitari Vall d'Hebron, Universitat Autònoma de Barcelona, Barcelona, Spain Hugo Vrenken: Department of Radiology and Nuclear Medicine, MS Center Amsterdam, Amsterdam, The Netherlands

Tarek Yousry: NMR Research Unit, Institute of Neurology, University College London, London, UK

Claudio Gasperini: Department of Neurology and Psychiatry, University of Rome Sapienza, Rome, Italy

